# Commercial influenza vaccines vary in both the structural arrangements of HA complexes and in induction of antibodies to cross-reactive HA epitopes

**DOI:** 10.1101/2022.09.22.509041

**Authors:** Mallory L. Myers, John R. Gallagher, Alexander J. Kim, Walker H. Payne, Kevin W. Bock, Udana Torian, Ian N. Moore, Audray K. Harris

## Abstract

Influenza virus infects millions of people annually and can cause global pandemics. Hemagglutinin (HA) is the primary component of commercial influenza vaccines (CIV), and antibody to HA is a primary correlate of protection. Persistent antigenic variation of HA requires that CIV be reformulated for new strains yearly. Differences in structural organization of HA has not been correlated with induction of broadly reactive antibodies, and CIV formulations can vary in how HA is organized. Using electron microscopy to study four current CIV, we found that these different formulations contained a variety of structures including: individual HAs, starfish-like structures with up to 12 HA molecules, and novel “spiked nanodisc” structures that displayed over 50 HA molecules along the complex’s perimeter. These spiked nanodiscs uniquely exposed conserved stem epitopes and elicited the highest levels of heterosubtypic cross-reactive antibodies. Overall, we found that HA structural organization can be an important CIV parameter and can be associated with the induction of cross-reactive antibodies to conserved HA epitopes.

## Introduction

Influenza is a widely prevalent respiratory virus with pandemic potential. Healthy individuals, even those who are vaccinated, are often still susceptible to influenza infection and may become sick, disrupting work and well-being ^1^. At-risk individuals that may be immunologically suppressed due to age or specific health issues can develop life threatening infections ^2^. When new zoonotic strains are created by reassortment between animal and human lineages, antigenically novel influenza virus puts even the healthy population at great risk, as observed in past influenza pandemics ^3^. Influenza antigenic variation has caused the commercial vaccine market for influenza to become a billion dollar industry ^4^.

Influenza is an enveloped virus, presenting 3 antigens on the viral surface. Hemagglutinin (HA), the most prevalent viral glycoprotein, is responsible for binding to target cell glycans, as well as execution of membrane fusion. Neuraminidase (NA) cleaves cellular glycans to assist in viral release from host cells, counteracting the binding activity of HA. Matrix (M2) is an ion channel embedded in the viral envelope which is integral for pH homeostasis of the virus ^5-7^. Following both natural infection with influenza virus and immunization with HA based vaccines, the majority of protective antibodies are targeted to HA ^8-10^. HA is a homotrimer consisting of three copies of the protein HA0 which can then be cleaved into two fragments, HA1 containing epitopes in the globular head region, and HA2 containing epitopes in the stem region of the glycoprotein. The highly conserved sequence of HA that executes fusion is in the HA2 stem region. Meanwhile, the HA1 head region, being most exposed on the virus surface, is both immunodominant and has a fast rate of sequence evolution. Yearly reformulation of influenza vaccines is required to ensure that HA antigenic variation does not escape vaccine-elicited antibodies ^11,12^.

Historically, influenza vaccine development was evaluated by antigenicity and protection in animal models focusing on the concentration of HA, but recently other viral components like NA and M2 are being explored ^13-15^. Commercially available influenza vaccines have been widely accessible in the United States since the 1950s ^16^ with the CDC regulating the concentration of HA in those offered ^8^. Early vaccine formulation strategies and procedures, many of which are still in use, required generation of antigen via seed viruses that are then used to infect embryonated chicken eggs, harvest virus, and extract HA by methods of varying specificity ^17^. To avoid the problem of egg-adapted HA mutations, some newer influenza vaccine designs are synthesized from influenza viruses grown in mammalian cells ^18^. Further developments in vaccine manufacturing were seen with the approval of an MF59 adjuvanted commercial vaccine ^19^, while another vaccine formulation utilized recombinant HA derived from a baculovirus expression system in insect cells ^20-22^. To date, structural characterization of commercial HA subunit vaccines for influenza has not extended beyond negative-stain electron microscopy (EM), without further image analysis ^23^. Those studies showed HA isolated from inactivated virus or produced by recombinant baculovirus expression presented rosette formations ^21,23,24^. Rosettes were naturally formed by removing lipid components, resulting in the transmembrane domain of multiple HA proteins being constricted into small lipid micelles. Within rosettes, the accessibility of the HA head domain was unperturbed, causing no concern for decreased immunogenicity as vaccine development had focused on the antigenically dominant head domain of HA. However, recent universal flu vaccine initiatives have shifted attention from a singular focus on HA head-targeted antibodies toward consideration of HA stem epitopes.

Pursuit of novel HA stem vaccines aiming to protect against both current and future influenza strains ignited interest in targeting the conserved stem of HA, which lies within the HA2 region near to the membrane and contains residues essential for viral fusion ^25,26^. At present, differences in the extent that existing influenza vaccine formulations display influenza stem epitopes has not been fully characterized. While some formulations offer better cross-protection against influenza strains that are not present in the vaccine, the mechanism for this effect in terms of HA organization and conserved epitope display is unknown ^19,27^. Addition of an adjuvant to a vaccine is known to directly stimulate the immune response in additional ways to subunit vaccines ^28^; however, adjuvant can also profoundly affect the higher-order organization of antigen in solution.

Here we explore the organization of HA within four licensed influenza vaccines (Fluzone, Flucelvax, Flublok, Fluad) by EM to identify structural correlates to the induction of antibodies against different influenza HA subtypes. We found that the four commercial influenza vaccines had HA components organized in heterogenous polymorphic complexes when examined by 2D imaging and classification by negative stain EM. While different in organization, all vaccines displayed the conserved stem HA epitopes and bound broadly reactive human anti-stem antibodies FI6v3 and CR6261. However, the four vaccines had important differences in their ability to elicit cross-reactive antibodies to HA subtypes not part of their formulation. All of the vaccines protected against a homologous H1N1 challenge, while only one vaccine, Fluad, elicited significant levels of antibodies to the stem and for HA subtypes not present in the vaccine formulation. This vaccine also presented a novel spiked nanodisc HA structure. Analysis of the spike nanodiscs indicated that HA stem epitopes were more readily accessible, suggesting a structural basis for the ability to induce heterospecific antibodies to subtypes not present in the vaccine. Induction of heterospecific antibodies will be important for the development of both more efficacious seasonal and universal influenza vaccines that elicit broad immune responses to antigenically divergent HAs.

## Results

### HA organization in different commercial vaccines

Negative-stain EM was used to determine if antigen structure and organization within the four commercial influenza vaccines were different. Of the four commercial vaccines studied, two were trivalent (Fluad, Fluzone), containing HA antigens for influenza H1 and H3 subtypes for influenza A, and HA Victoria lineage for influenza B. The quadrivalent vaccines (Flublok, Flucelvax) had an additional influenza B HA Yamagata lineage added (Fig. S1). All vaccine HA antigens were generated from inactivated influenza virus except for the Flublok vaccine, which was composed of recombinant HA protein (Fig. S1). Western blot analyses of the vaccines’ components confirmed the presence of the major antigens of the commercial vaccines (Fig. S2A, S2B, S2C), including HA proteins of H1 and H3 subtypes in addition to influenza B HA glycoproteins. Also confirmed present were other viral components detected at lower levels and exclusively in virus-derived formulations. These other viral components included neuraminidase, matrix, and nucleoprotein (Fig. S2D, S2E, S2F).

Four configurations summarized the organization of HA in these vaccine formulations: (1) isolated HA molecules, (2) starfish-like complexes with HA radiating from a central core, (3) HA in micelles, and (4) HA in a larger disc-like structure (Fig.1). In EM micrographs, glycoproteins appear as a lighter shade of gray against a darker background, since the negative stain is excluded by presence of the protein. On rare occurrences, genomic RNP complexes were also visible (Fig 1A, asterisk). Class-averages were generated from thousands of picked particles in order to visualize the most common structures observed across the entire sample. In general, average structures indicated that the conformation of HA was presented in the characteristic peanut-shapes of prefusion trimeric molecules, which were in multimeric starfish-like complexes (e.g. Fig. 1B, left, middle panels) and as isolated HA trimers associated with small volumes of membrane (e.g. Fig. 1B, right panel).

**Fig. 1.**
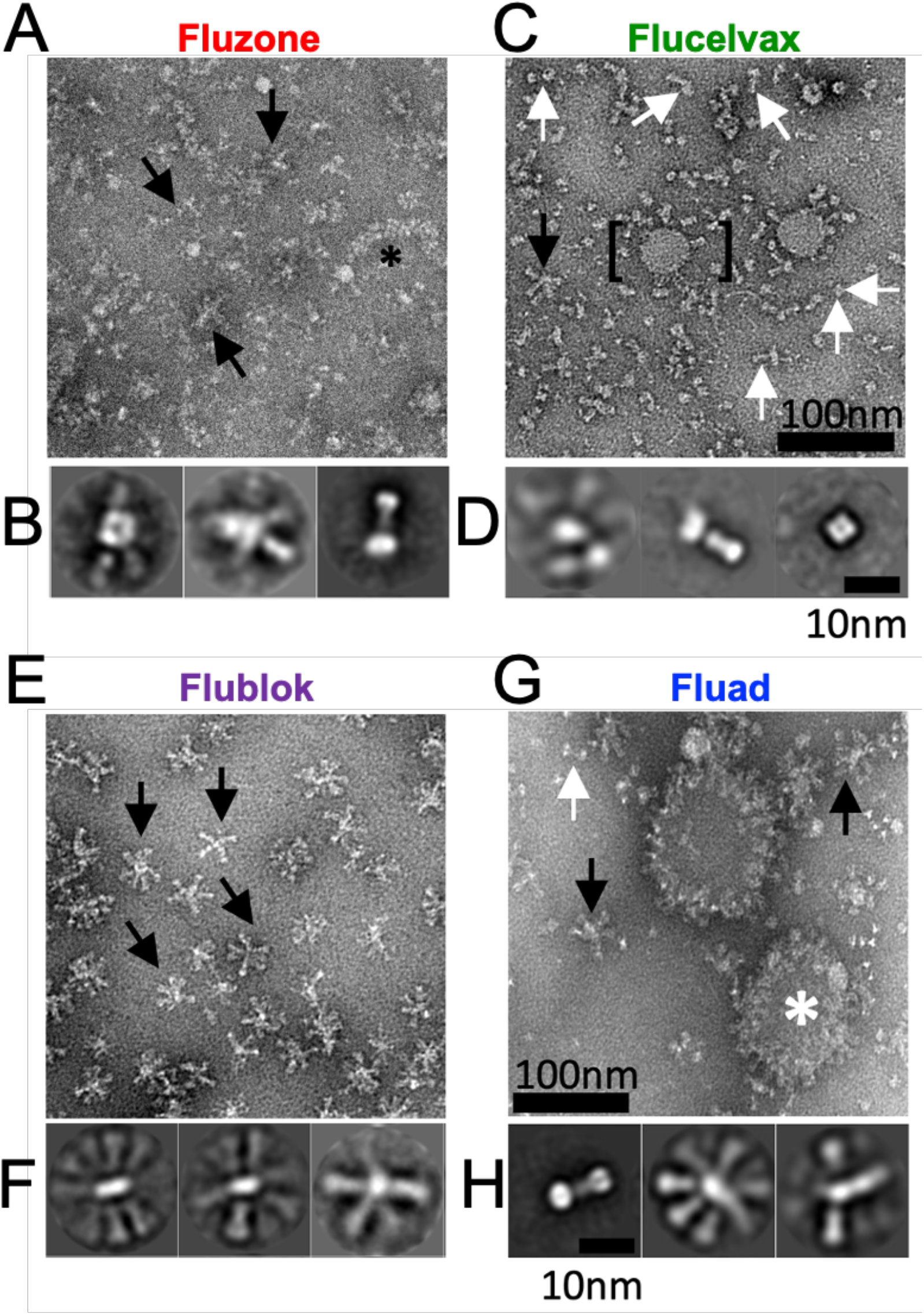
Structural organization of HA antigens in influenza vaccines. (A) Images from negative-stain EM of Fluzone vaccine and (B) select 2D classes averages. Similarly images and class averages for (C,D) Flucelvax, (E,F) Flublok, and (G,H) Fluad influenza vaccines. Complexes observed include HA starfish-like complexes (black arrows), isolated molecules (white arrows), large round complexes (white asterisk), RNP (black asterisk) and small round mostly bald particles (bracket, putative micelles).

Starfish-like structures were characterized by glycoprotein spikes protruding from a central location containing HA transmembrane domains. Starfish-like complexes of HA and isolated HA trimers were present in Fluzone (Fig. 1A, black arrows). Flucelvax also contained both isolated HAs and starfish complexes, but differed from Fluzone in that isolated HA molecules were more prevalent (Fig.1C, white arrows) than starfish-like complexes (Fig. 1C, black arrow). Class averages of Flucelvax indicated the presence of HA-starfish-like complexes, isolated HA molecules, and NA heads which were identifiable by their tetrameric symmetry (Fig. 1D). Unique to Flucelvax were relatively large 100-200 nm lipid structures (i.e. micelles) with sparsely distributed HA proteins protruding from their surface (Fig. 1C, black bracket). Flublok was the most structurally homogeneous, with HA organized exclusively in starfish-like complexes, commonly containing 5-12 HA trimers (Fig. 1E). Constituent HA molecules presented a prefusion peanut-shape, similar to the HA starfish structures presented in the class-averages of Flublok (Fig. 1F). As demonstrated by class-averages, Fluad contained isolated HA molecules and HA-starfish complexes similar to those found in other vaccines (Figs. 1G and 1H). However, Fluad contained an additional component not seen in other formulations. Large round complexes were observed with glycoprotein spikes dotting their outer perimeter (Fig. 1G, white asterisk). Many HA proteins on these complexes were visible as top-views with the HA spikes oriented towards at the camera, sometimes permitting the triangular organization of the trimer to be observed. Different years of each of these influenza subunit vaccines were compared and the structures for each vaccine matched the EM data of figure 1. Thus, confirmatory EM results indicated that the organization of HA molecules was consistent from year to year (data not shown).

### Modeling the effects of HA stem epitope crowding

The observed differences in HA molecule spacing and arrangement across different vaccine complexes led us to investigate the likelihood that HA stem epitopes were blocked or occluded by steric hindrance from neighboring HA molecules. We enumerated possible pairwise HA orientations by Monte Carlo simulation and evaluated if these configurations would occlude stem antibody binding (Fig. 2A, 2B). Test configurations started with the atomic coordinates of a trimeric HA ectodomain with all three stem epitope binding sites occupied by fragment antigen-binding (Fab). Then a second HA-trimer with bound Fabs was placed randomly, proximal to the first, and the configuration was assessed for steric clash (Fig. 2A). The random placement of pairs of HA-Fab complexes and their evaluation for steric clash was a Monte Carlo method to empirically sample the irregular parameter space relating HA distances, relative angles, and steric clash (Fig. 2B). We segmented the resulting plot of configurations of HA-Fab complexes into three regions: (i) HA-Fab complexes always clashed with each other, (ii) stem Fabs interdigitated and may or may not clash depending on the HA rotation and (iii) HA-Fabs never clashed due to insufficient spacing that no axial rotation of HA could alleviate the steric clash (Fig. 2B). The Monte Carlo simulation sampled 500,000 random conformations, which empirically enumerated the steric clash landscape used to define these three regions. The orientations generated by the Monte Carlo method describe the configuration space possible between proximal HA trimers in starfish-like presentations that were observed by EM. At certain angles between HA-trimers, clashes may occur between stem-Fabs even when the inter-HA distance is comparatively large (16 nm) relative to the Fab domain size (7 nm), effectively reducing the Fab binding in those circumstances. The simulations illustrated how increased angles between the long axis of HA molecules allow full occupancy of stem epitopes at shorter distances between HA trimers (Fig. 2C vs 2D). Thus, smaller diameter complexes with greater curvature more readily exposed their full set of stem epitopes to antibody binding, partially counteracting the presence of fewer antigens on these smaller starfish complexes.

**Fig. 2.**
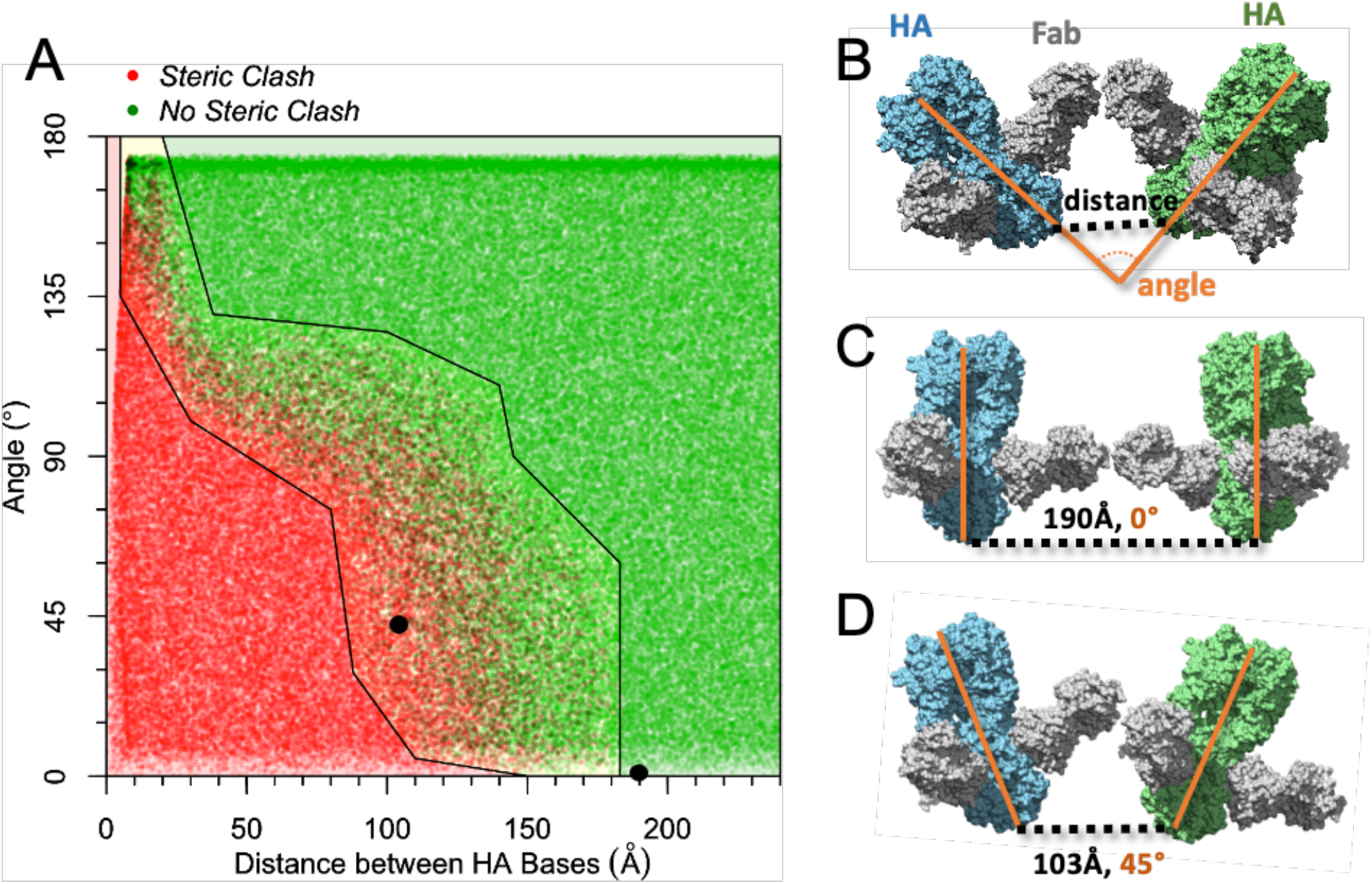
Monte Carlo model relating HA orientation with occlusion of stem antibody binding. (A). Structures of pairs of HA-trimers, each with 3 stem Fabs bound in complex, were randomly oriented by rigid rotation relative to each other, and then the resulting structures were scored for steric clash. The steric clashes were tabulated according to the distance between the base of HA trimers, and the angle between the central axis of HA trimers. The plot of steric clashes was empirically segmented into 3 regions: no steric clash, all steric clash, and conditional steric clash depending upon the interdigitation of Fabs. Within the central region of conditional steric clash, 57% of these configurations free of steric clash. The two black points in panel A represent the example complexes depicted in panels C and D. (B) A visualization of the steric clash model system denotes the distance between HA bases, and the angle between central axes of HA trimers. (C) An example configuration shows parallel HA trimers (angle = 0°) and Fab domains rotated to face each other, but the 190 Å distance between bases avoids steric clash. (D) An example of HA trimers in the conditional clash region where Fabs interdigitate, avoiding clash in some rotations, but Fabs could easily clash in other rotations of the HA trimer.

### 3D structure of HA starfish in Flublok

To determine the structural organization of starfish-like complexes, we used cryo-electron tomography of HA complexes in Flublok (Fig. 3). Virtual slices through Flublok complexes in the 3D tomogram revealed that HA trimers radiated out from a central location in 3D space (Fig. 3A-3F, Movie S1). While the technique of negative staining had flattened the HA-starfish complexes during visualization, cryo-electron microscope revealed that the solution structure of HA complexes indeed emanated outward from a central clustering of HA transmembrane domains (Fig. 3G, 3H). A schematic model in which trimeric HA molecules cluster into a starfish complex via their transmembrane domains forming a central hydrophobic core region is depicted (Fig. 3I, grey core). Some starfish complexes were elongated rather than spherical, with HAs emanating from a cylindrical lipid core rather than a single point. The average diameter of HA complexes was approximately 25 nm. The density profiles of the constituent HA molecules were consistent with the prefusion state of HA, one which is competent for stem-epitope binding by HA-stem targeted antibodies.

**Fig. 3.**
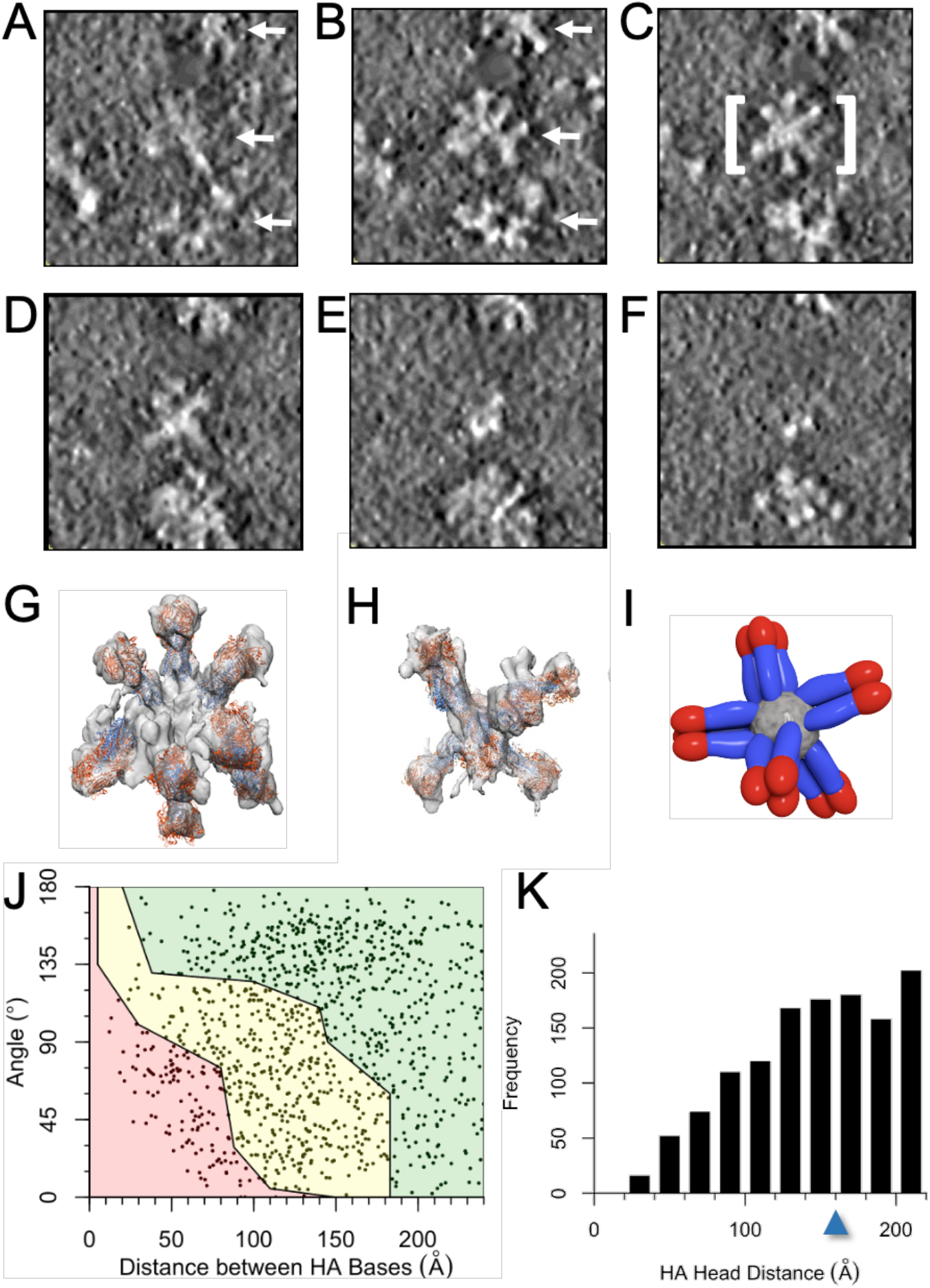
Flublok structural analysis by cryo-electron tomography and molecular modeling. (A-F) Example of serial slices from top to bottom of a cryo-electron tomogram with a field of HA complexes (white arrows). (A, B) White arrows in panels A and B indicate the location of three individual starfish-like complexes made of constituent HA molecules. (C) View in which the slice of the central complex (white brackets) presents a starfish like arrangement of its HA molecules. (G) a molecular model of the complex with trimeric HA ectodomain coordinates docked into the 3D tomogram shown as a transparent isosurface. (H) A different starfish-like complex is displayed with HA trimers docked into the tomographic map, shown as a transparent isosurface. (I) A schematic of a HA-starfish structure illustrates how a central hydrophobic core (gray) contains HA transmembrane domains and lipid, and HA proteins point outward exposing the HA head colored in red, and the HA stem colored in blue. (J) Distance and angular measurements between HA-trimer bases, according to HA ectodomains docked into tomograms of the HA starfish-like complexes. The plot is segmented into colored regions according to Monte Carlo simulations of HA-stem clashes. (K) HA-trimer distances between the central axis of head domains in docked complexes were tabulated and plotted as a histogram. The arrow denotes the threshold distance of 16 nm under which bivalent binding is possible.

HA-trimers were modeled into each density with dimensions appropriate for HA (Fig. 3G, 3H), which allowed a series of HA starfish complexes from cryo-electron tomography to be characterized for steric hinderance of stem epitope binding based upon the metrics established by Monte Carlo simulation. By plotting these experimental HA-trimer configurations on the same landscape enumerated previously (Fig. 2A), the probability of stem epitope availability was estimated. For Flublok, there was a broad distribution of angles and distances between HA molecules (Fig. 3J). The likelihood that the HA configuration in Flublok permitted binding to a complete complement of stem antibodies was estimated as 100% of HAs in the permissive region of the Monte Carlo simulation plus the HAs in the conditional clash normalized by the fraction estimated competent for full stem antibody binding by Monte Carlo simulation (57%). For Flublok, 49% of HAs were in the permissive region and 40% were in the conditionally permissive region. After scaling the fraction in the conditionally permissive region by 57%, a final estimate for the fraction of stem epitopes fully capable of binding stem antibodies in Flublok was 72%. Configuration of HAs can also impact the propensity for bivalent binding by a single antibody to two different HA heads. Bivalent binding effectively increases the affinity of the antibody for the epitope by decreasing the off rate, enabling weak but conserved epitopes to progress through affinity maturation. The threshold for bivalent binding has been experimentally determined as approximately 16 nm ^29^. Influenza vaccines that orient HA heads within 16 nm have the potential for bivalent binding, and this potential is visualized as the frequency of HA heads below this distance threshold (Fig. 3K). 63% of HA pairs in Flublok within 22 nm are within this threshold for bivalent binding.

### 3D structure of HA spiked nanodiscs in Fluad

Cryo-electron tomography of Fluad was performed to probe the 3D molecular organization and spatial display of HA stem epitopes. Fluad is a vaccine adjuvanted with MF59 (squalene oil-in water emulsion), and we aimed to visualize the effect of the adjuvant. Virtual slices through Fluad tomograms revealed round vesicle-like structures interpreted to be the MF59 adjuvant (Fig. 4A, 4B, asterisks). Interspersed among the vesicles were flat disc-like structures approximately 100 nm in width, containing glycoprotein spikes (Fig. 4A, 4B, arrows, Movie S2). We refer to these structures as “spiked nanodiscs”. HA ectodomains could be modeled into the spike densities at the periphery of the spiked nanodiscs (Fig. 4C, 4D). In contrast to Flublok, the Fluad 100 nm spiked nanodiscs are larger and can contain over 50 HA trimers. To observe the nature of the lipid disc constituent in the spiked nanodiscs, we performed subtomogram averaging from 58 discs. Averaging just the center of the discs where no HA was present, we measured a distance of 48 Å between two peaks of density in a cross-section (Fig. 4E). We concluded that the spiked nanodiscs were comprised of a lipid bilayer (Fig. 4E). HA glycoprotein spikes studded the outer perimeter of the spiked nanodisc, oriented normal to the flat surface of the spike nanodisc. Thus, pairs of HA trimers were approximately parallel when located on the same side of the spiked nanodisc, and antiparallel when on opposite sides of the spiked nanodisc (Fig. 4F, Movie S3). This exposed the full height of HA at the edge of the spiked nanodiscs, and prominently displayed conserved stem epitopes. The fraction of HAs in the spiked nanodiscs that were permissive to full stem-antibody binding was estimated as the sum of HAs in the permissive region (67%) and the HAs in the conditionally permissive region (30%) scaled by 57% (fraction available from Monte Caro simulations). Therefore, the final estimate for the fraction of HAs in Fluad permissive to full stem antibody binding was 84% (Fig. 4G). Bivalent binding, approximated by the frequency of HA heads being within 16 nm, was estimated at 62% (Fig. 4H) within the locale of 22 nm, notably similar to bivalent binding permissibility of Flublok starfish.

**Fig. 4.**
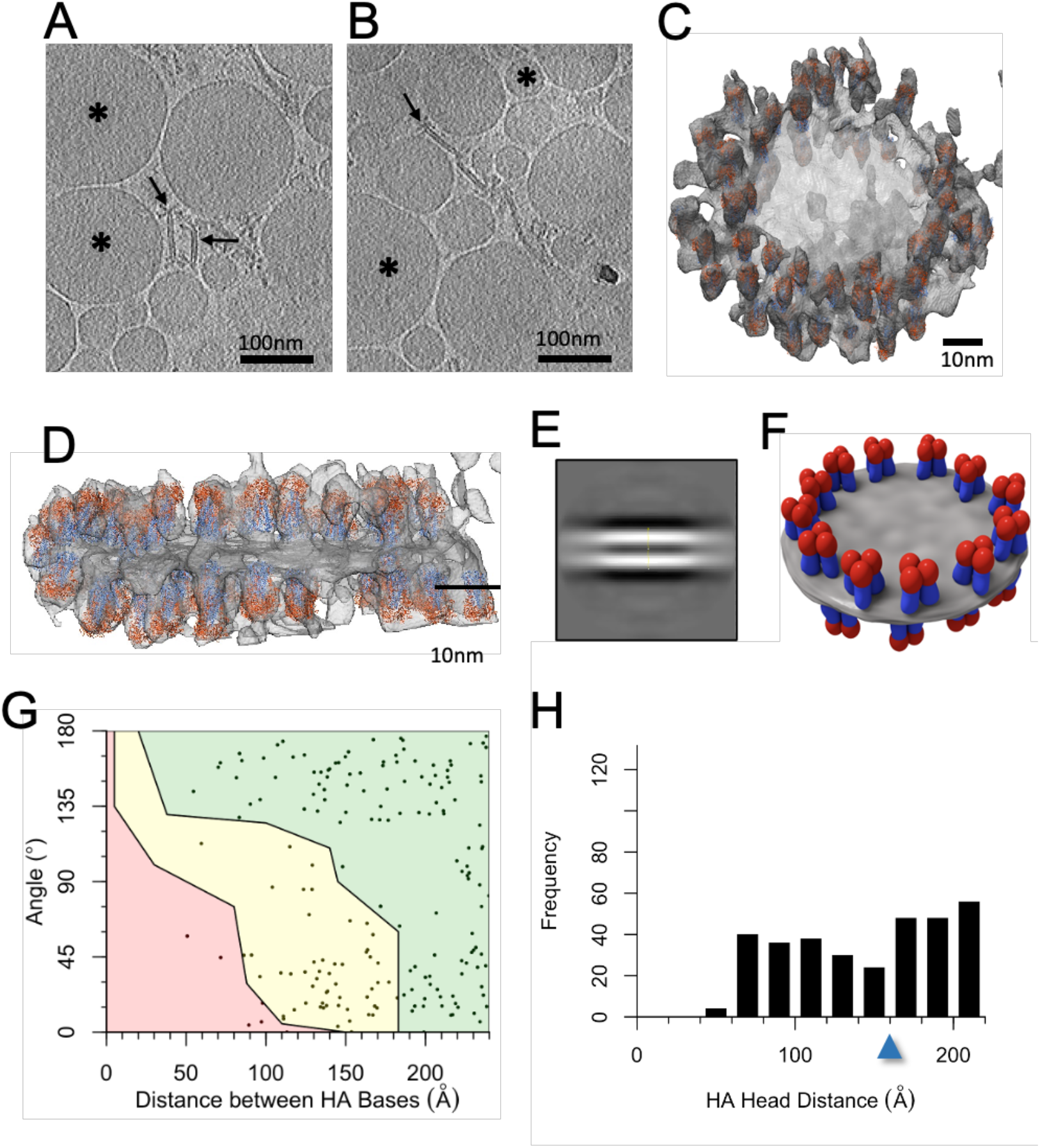
Fluad structural analysis by cryo-electron tomography and molecular modeling. (A, B) Examples of slices through cryo-electron tomograms with a field of Fluad spike complexes and putative adjuvant vesicles. Black arrows indicate some spike complexes protruding from an apparent lipid bilayer. Black asterisks denote adjuvant vesicles. (C) Oblique view of a cryoelectron-tomographic 3D reconstruction (gray) of a Fluad large particle. A flat disc-like structure contains spiked densities around its perimeter, which were used to dock HA molecules (ectodomain trimers, PDB ID 3LZG) found to be approximately parallel and anti-parallel on either side of the disk. (D) Side-view of the Fluad particle in panel C illustrates parallel and anti-parallel HA orientation. (E) Side view of a subtomogram average of Fluad disc-like structures, focusing on the disc center. Two layers of density are visible in a density slice bisecting the disc, which indicated the discs are composed of a bilayer. Distance between the centers of each band of density were 48 Å. (F) A schematic of the HA trimers on the perimeter of the lipid-based disc (gray) illustrates their consensus orientations with HA heads, colored in red, aligned normal to the disc surface. HA stem regions are colored in blue. (G) Distance and angular measurements between HA-trimer bases of HAs docked into a series of Fluad discs. The plot is segmented into colored regions according to Monte Carlo simulations of HA-stem clashes. (H). HA-trimer distances between the central axis of head domains were tabulated and plotted as a histogram. The arrow denotes the threshold distance of 16 nm under which bivalent binding is possible.

### HA stem display and induction of heterologous HA subtype antibodies

Differences in HA stem epitope display from structural studies of commercial influenza vaccines led us to more closely investigate the *in vivo* immunogenicity of these same formulations. To probe these differences, we assessed binding (*in vitro* display) by ELISA using stem antibodies and recombinant proteins. Anti-stem monoclonal antibodies FI6v3, CR6261, and C179 were employed to probe differences in the availability of stem binding epitopes in the commercial influenza vaccines Fluad, Flublok, Fluzone, and Flucelvax (Fig. 5A). All vaccines contained HA protein with binding epitopes for anti-HA stem antibodies FI6v3 and CR6261. However, after normalizing for HA concentration in different formulations, area under the curve analysis indicated that both Flublok and Fluzone had more stem epitope binding availability than Fluad and Flucelvax (Fig. 5A). These results indicated that the binding availability of conserved HA stem epitopes could vary between different commercial influenza vaccines that were purified and formulated by different methods, despite their similar composition of full-length HA trimers (Fig. S1).

**Fig. 5.**
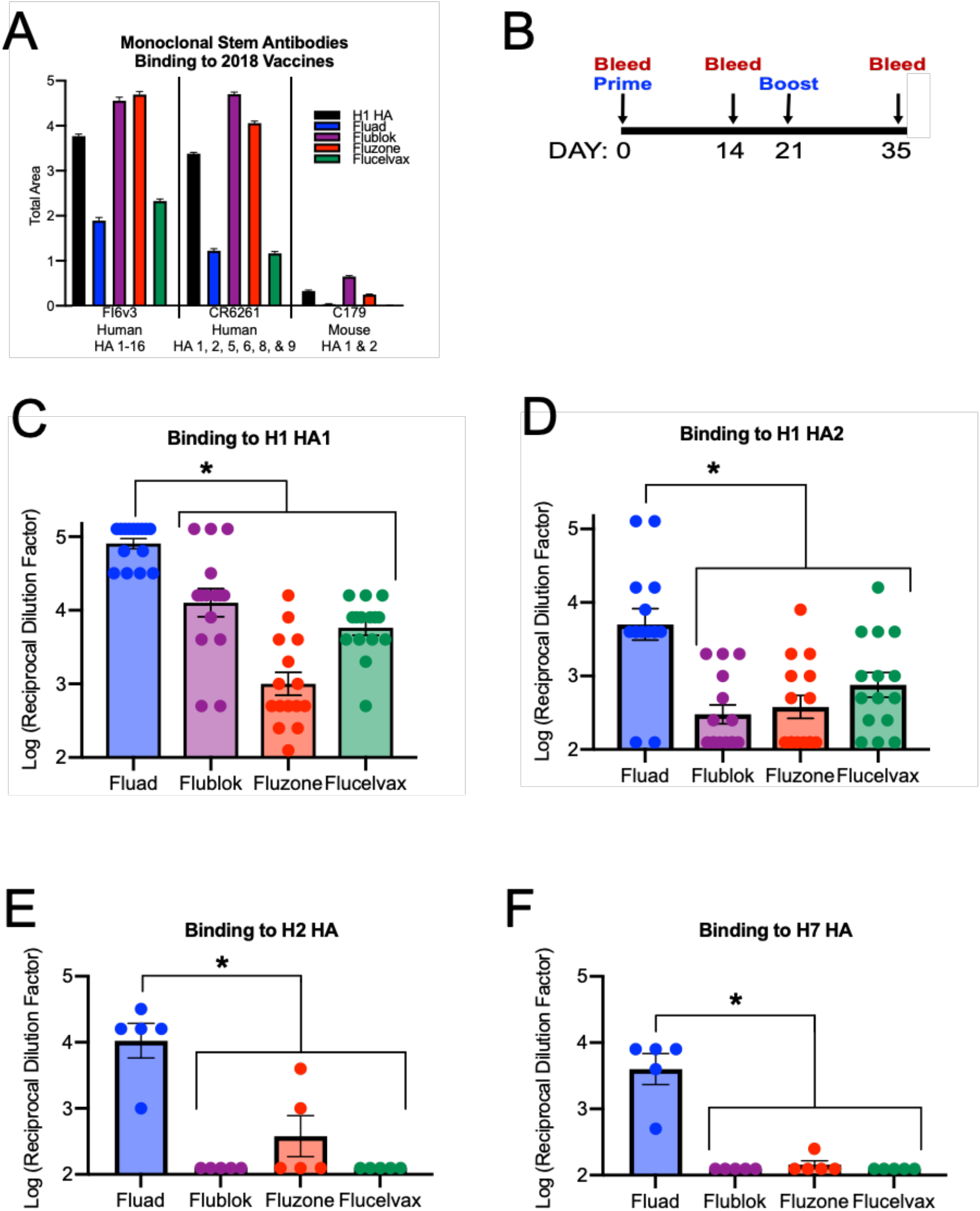
Commercial vaccine stem-epitope availability and cross-reactive immunogenicity. (A) ELISA binding analysis of commercial vaccines (Fluad, Flublok, Fluzone, Flucelvax) by HA stem monoclonal antibodies (FI6v3, CR6261, C179). Vaccine samples were normalized for H1 HA amount prior to plating. Recombinant H1 HA protein (H1 HA A/Michigan/45/21015, black bars) was a positive control. (B) Schedule for immunization of mice with vaccines with day 35 sera used in subsequent ELISA studies. (C-F) The endpoint titer levels of different vaccine sera were compared for cross-reactivity to different recombinant proteins: (C) H1 HA1 head, (D) H1 HA2 stem, (E) H2 HA, and (F) H7 HA. PBS control sera was also tested to determine threshold, but not shown.

*In vivo* immunogenicity was tested to observe if *in vitro* stem epitope display would be predictive of stem antibodies elicited *in vivo* by the commercial vaccines. Mice were immunized with the vaccines using a prime and boost immunization schedule (Fig. 5B). Vaccine HA antigens were immunogenic and elicited homosubtypic binding antibodies to their H1, H3, and influenza B HA antigens, as judged by ELISAs using immunized mouse sera and full-length recombinant HA proteins (Fig. S3). When comparing endpoint titers calculated from dilution curves (Fig. S4 and Fig. S5) there were statistically significant differences between the different vaccine sera binding to H1 HA1 head, H1 stem, H2 HA and H7 HA recombinant proteins (Fig. 5C-5F). Fluad sera had higher levels of antibodies binding to a H1 HA1 head domain protein than the sera from the other three vaccines (Flublok, Fluzone, Flucelvax) (Fig. 5C). Likewise, Fluad elicited higher levels of cross-reactive antibodies to an H1 stem protein (Fig. 5D) when compared than the other vaccines. Although H2 and H7 HA antigens were not components of the vaccines, Fluad elicited statistically significant higher levels of cross-reactive antibodies to both H2 HA and H7 HA when compared to the other vaccines, indicating the elicitation of higher levels of heterosubtypic antibodies by Fluad (Fig. 5E-F).

### Weight-loss and lung pathology differences between vaccines

Having observed differences in both the vaccines’ organizations and the elicitation of cross-reactive antibodies, we challenged mice with a homotypic H1N1 viral challenge and looked for differences in survival, weight loss (Fig. 6), and lung pathology (Fig. 7). As expected, all vaccines protected mice from H1N1 challenge (Fig. 6B). This was expected because all the vaccines contained H1 HA protein and elicited antibodies to H1 HA (Fig. S1-S3). However, further examination of the weight-loss curves showed that Fluad and Flublok had trends for reduced weight loss compared to both Flucelvax and Fluzone (Fig. 6C). The most observable difference was at day 3 (Fig. 6C, arrow). When the experimental timeline was repeated with additional mice for the collection of lung tissue, the weight-loss differences observed on day 3 post challenge were statistically significant (Fig. 6D). Vaccination with any vaccine followed by challenge resulted in lesser weight loss when compared to the saline control group. Between the vaccine groups, mice given Flublok and Fluad had no significant weight-loss difference between each other after challenge but did have a statistically significant reduced weight-loss when compared to Fluzone and Flucelvax (Fig. 6D). Interestingly, when lung tissue was homogenized, infectious virus could not be recovered from the lungs of mice vaccinated with Fluad, while the other vaccines permitted viral recovery (50% Tissue Culture Infectious Dose (TCID50) assays) (Fig. 6E). Although Fluad and Flublok did not show a statistical difference in weight-loss reduction between them, they differed in their ability to permit recovery of virus from the lungs (Fig. 6E).

**Fig. 6.**
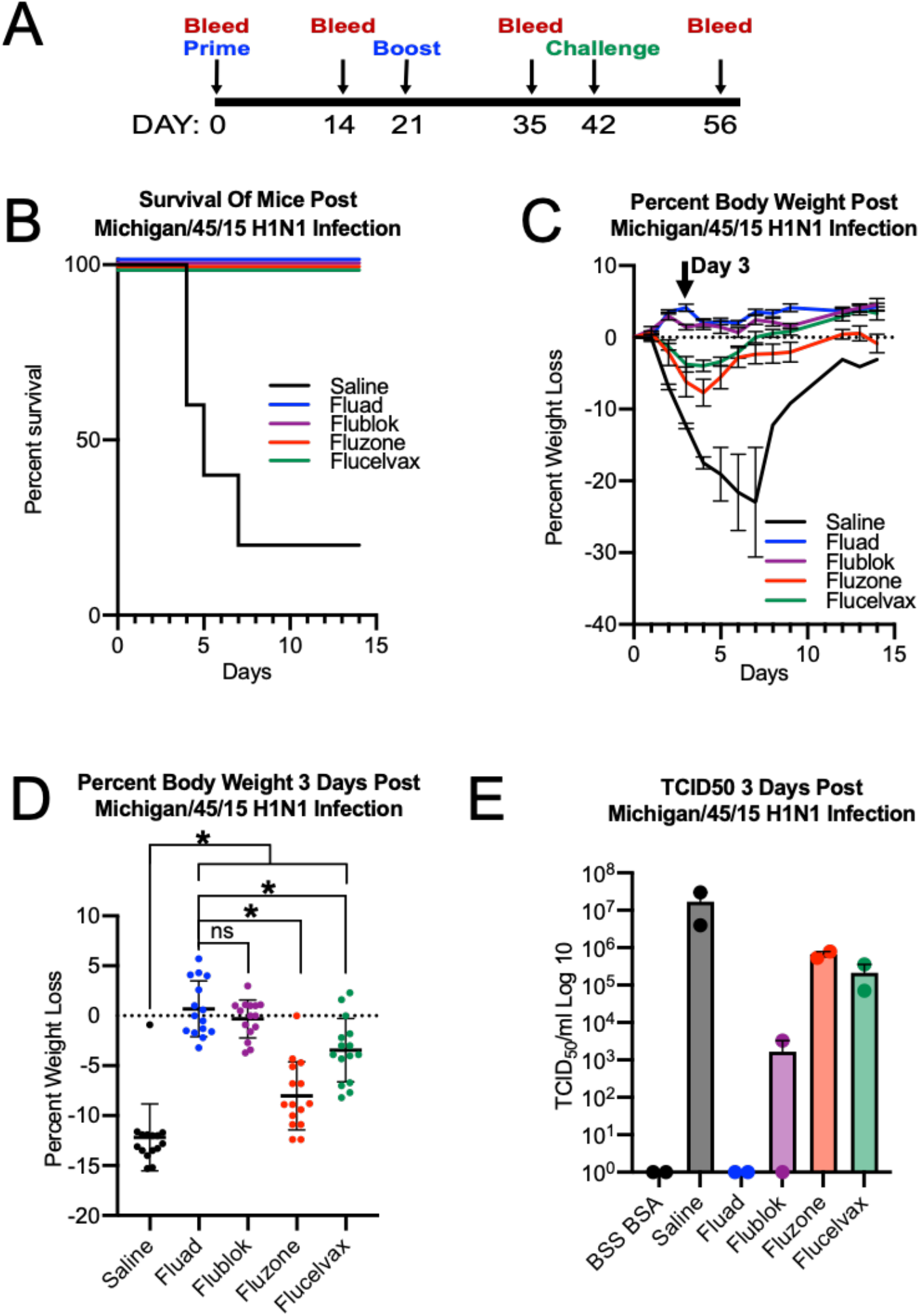
Protection efficacy and weight lost differences after H1N1 challenge. (A) Mice were immunized with commercial influenza vaccines (day 0 and 21) and challenge with A/Michigan/45/2015 H1N1 influenza virus at 10x the lethal dose for control mice (day 42). (B) Survival curves for mice immunized with saline control (black), Fluad (blue), Flublok (purple), Fluzone (red) and Flucelvax (green). (C) Weight-loss curves for challenged mice that were immunized with saline (black), Fluad (blue), Flublok (purple), Fluzone (red) and Flucelvax (green). The black arrow indicates weight-loss on day 3 post H1N1 challenge. (D) Statistical analysis of weight-loss differences (N=15 mice) between immunization groups for day 3 post H1N1 challenge. No statistical difference in weight-loss between Flublok and Fluad is denoted (i.e., ns.). Asterisks denote statistical differences (*P*<0.05) in weight-loss between saline control as compared to vaccines (Flublok, Fluad, Fluzone, Flucelvax) and statistical differences between Flublok and Fluad versus the other vaccines, Fluzone and Flucelvax. (E) Mouse lung titers were determined by 50% Tissue Culture infectious Dose (TCID50) for tissues taken day 3 post-H1N1 challenge and compared between commercial influenza vaccines.

**Fig. 7.**
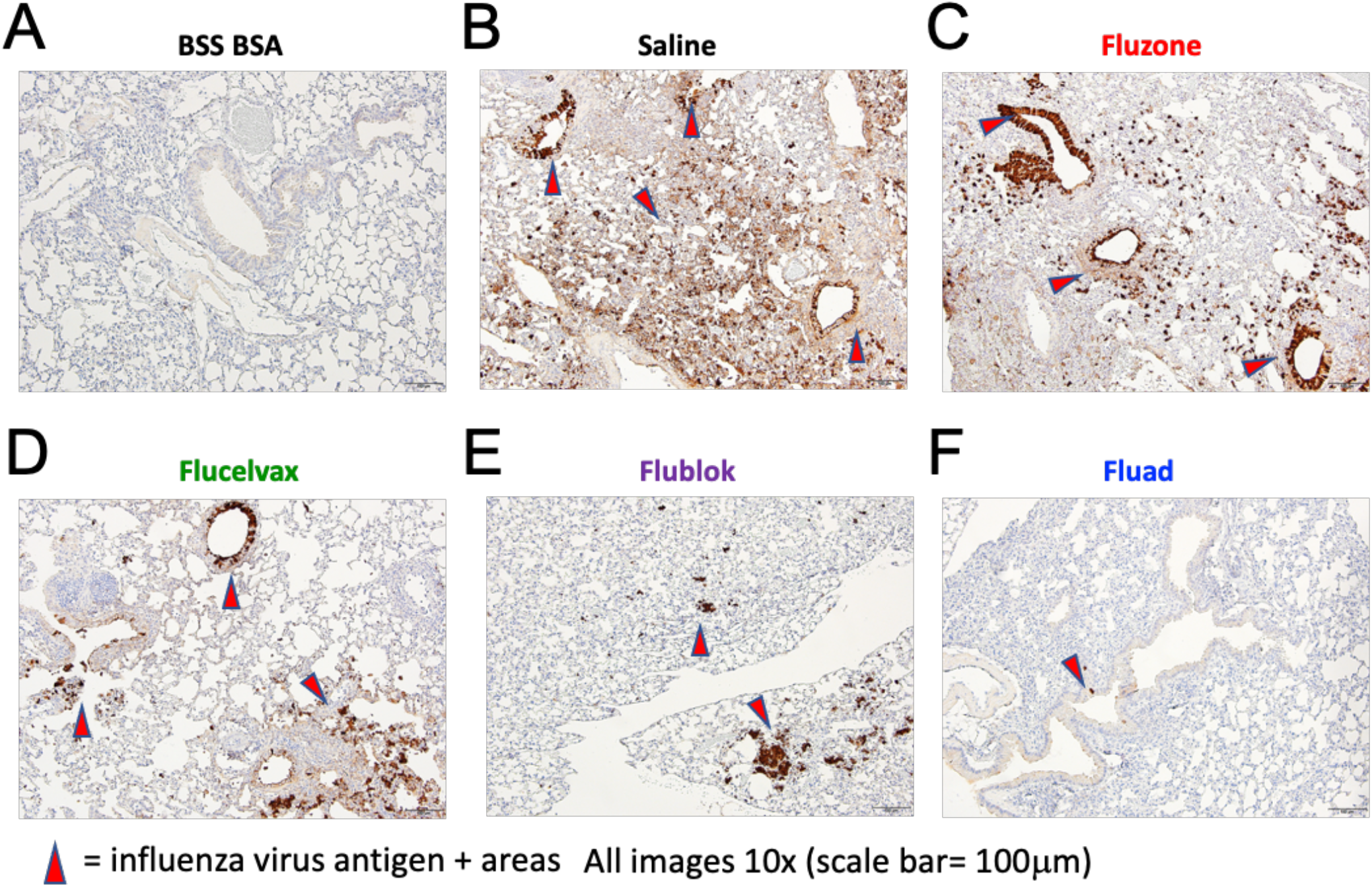
Histopathology in mice lungs following vaccination with commercial influenza vaccines and subsequent challenge with H1N1 influenza virus. (A-F) Images show immunohistochemistry (IHC) against the influenza nucleoprotein protein for different vaccination groups. Images shown are from day 3 post H1N1 challenge for all groups. (A) Healthy lung control via immunization with saline and challenged with saline-BSA. (B) Pathogenic positive control via immunization with saline with challenge with H1N1 virus. (C-F) Immunization with the commercial influenza vaccines (C) Fluzone, (D) Flucelvax, (E) Flublok, and (F) Fluad and subsequent challenge with H1N1 virus. All images at 10x, (scale bar is 100μm). Red arrows denote viral antigen areas.

Lung samples were also examined by immunohistochemistry at day 3 post-viral challenge. An unbiased, blinded method was employed for analysis of histopathology of mouse lungs following H1N1 challenge (Fig. S6). Processed samples indicated that when compared to the healthy-lung control (Fig. 7A), mice receiving saline instead of vaccine had large amounts of influenza NP antigen staining in lung sections (Fig 7B, red arrows). Mice vaccinated with Fluzone and Flucelvax had similar distributions of viral antigen in the lungs. Animals given Flublok had lesser antigen distribution than Fluzone and Flucelvax. Fluad had the least amount of antigen detected in the lungs of all the vaccines tested. Fluad was visually the most similar to the healthy-lung control (Fig. 7).

## Discussion

Influenza hemagglutinin is a major target of the neutralizing antibody response to influenza, and antibodies elicited by the immunodominant HA head domain are evaded by antigenic drift in just a few influenza seasons ^30^. Antibodies targeting the HA stem are frequently broadly reactive across influenza subtypes but are elicited in low amounts by seasonal influenza ^31^. To understand if different vaccine formulations impact the nature of the antibody response and prevalence of cross-reactive stem-binding antibodies, we evaluated 4 subunit influenza vaccines that capture the varying methods of formulating HA for vaccination (i.e., Fluzone, Flucelvax, Flublok, Fluad) (Fig. 1, S1). Recent studies using structure-guided nanoparticle designs for viral vaccines have indicated numerous important factors for glycoprotein display and immunogenicity. These factors may include glycoprotein pre/post-fusion conformation, glycoprotein spacing, and number of glycoproteins or multivalency of the nanoparticle ^26,32,33^. However, these factors such as HA conformation, spacing and numbers have not been studied in terms of current influenza commercial vaccines in a side-by-side comparison. Other types of antigen display are currently being developed to enhance immunogenicity of vaccines, such as octahedral ferritin particles, to display the prefusion trimeric ectodomains of a multitude of different viral proteins such as influenza HA, HIV Env, and SARS-CoV-2 spike ^26,33-35^. Although commercial vaccines are given to millions of people world-wide, differences in stem epitope display and vaccine structure between a group of influenza vaccines have generally not been studied. Previous reports have been limited to biochemical or human/animal studies for individual commercial influenza vaccines ^19,36^.

In this work, we directly observe a variety of parameters that describe how epitopes are made available in commercial influenza vaccines. We started by quantitating epitope bioavailability via monoclonal antibody binding. By normalizing the concentration of H1 HA protein, we found that the highest fraction of H1 HA stem was bound in Flublok and Fluzone formulations, while Flucelvax and Fluad bound the least stem antibodies (Fig. 5A). We also found differences in the induction of stem-targeted antibodies in mice, which did not follow the same trend as available stem epitopes in vaccine formulations (Fig. 5D-F). While Fluzone and Flublok exposed the most stem epitopes *in vitro*, these two vaccines induced less HA2 (i.e. stem region) specific antibodies in mice. Meanwhile, Fluad exposed among the fewest stem epitopes *in vitro* while eliciting the most HA2 (i.e. stem region) specific antibodies in mice. Fluad elicited the most HA1-specific antibodies, targeting the HA immunodominant head. Fluzone contained the most non-HA viral components and offered the most heterogenous configurations of HA clusters, while inducing the least amount of HA1 head specific HA antibodies. We found that HA complexes in commercial influenza vaccines did not exhibit any regular symmetry (Fig. 1, 3, 4). Although some HA starfish-like complexes did have eight HA trimers like the engineered HA ferritin nanoparticles ^26,33^, commercial vaccines contained full-length HAs. Our structural studies suggests that in the vaccines, the HA molecules are in a prefusion state and that the mechanism of HA complex formation appears to be via aggregation of the hydrophobic transmembrane (TM) regions (Fig. 3G, 3H).

Molecular models from 3D cryo-electron tomography indicate that in Flublok, starfish-like structures are bound together by hydrophobic TM regions which partition out of aqueous solution (Fig. 3A-3I). In contrast, HA trimers in Fluad were arranged as a double ring with a disc shape that we refer to as a spiked nanodiscs (Fig. 4). This nanodisc structure is a different geometrical display of HA molecules than both the HA starfish arrangement and rigid nanoparticles, such as those scaffolded by ferritin ^26,33^.

The Fluad spiked nanodisc structure could be a novel method of displaying viral glycoproteins that is optimized to present conserved stem epitopes. These larger spiked nanodiscs also contain more HA trimers than most nanoparticle scaffolds, perhaps equivalent to icosahedral symmetry vehicles. Compared to other subunit vaccines, Fluad has higher multivalency of HA complexes (Fig. 4C vs 3G). While Fluad complexes would have less multivalency than whole virus or virosomes, the particle concentration of these virus-sized complexes would also be lower and would require direct comparison to evaluate.

Ideal spacing and presentation of antigen for immune stimulation has long been an interest when rationally designing vaccines ^32^. Densely spaced antigens on a particle surface offer more binding sites to interact with immune system components, yet very densely packed antigens can occlude access to epitopes not on the top of the antigen. Using a Monte Carlo method to evaluate the irregular molecular surfaces of HA trimers bound to stem-Fab, we found that HA stem epitopes are readily occluded when HA molecules pack closely next to each other, yet stem epitopes become more available as the angle between HA trimers increases (Fig. 2). Thus, there is an interesting relationship between multivalency and epitope accessibility, as increased density of HA antigen can both occlude epitopes and offer more opportunity for bivalent binding. Meanwhile, smaller complexes with lower multivalency gain epitope accessibility through less HA being present, as well as greater particle curvature. Previous studies with viral glycoproteins, such as Epstein-Barr virus gp350 and influenza HA trimer indicated that increasing multivalency via a ferritin nanoparticle yielded increased immunogenicity and vaccine efficacy in animal models ^33,37^. For influenza commercial vaccines, our work suggests that low multivalency could give higher HA stem *in vitro* binding activity by anti-HA stem monoclonals like for Fluzone (Fig. 5A red bars), but not elicit higher antibodies to the stem or cross-reactive to different HA subtypes *in vivo* (Fig. 5D-F). Fluzone contained HA starfish-like complexes of relatively low numbers of HA (i.e. low multivalency) (Fig. 1A, 1B). Almost paradoxically, Fluad had lesser *in vitro* binding by the human anti-stem broadly reactive antibody FI6v3 (Fig. 5A blue bars) yet elicited the highest levels of cross-reactive antibodies to the stem and to HA subtypes not constituent in the vaccines (Fig. 5D-5F). This ability of Fluad to elicit cross-reactive antibodies coincided with the observation by cryo-electron microscopy of the unique spiked nanodiscs, containing a variable number of HAs commonly over 50. The uniqueness of the spiked nanodisc structure in the display of parallel HA trimers exposing stem epitopes around the perimeter in a fashion that also promotes multivalent binding specifically of the stem epitope suggests the adjuvant may act to directly arrange the HA trimers to improve the elicitation of cross-reactive antibodies.

In conclusion, although all of the commercial influenza vaccines in this study contained full-length HA proteins there were differences in the number of HA molecules per particle (multivalency), spacing, and epitope display within HA complexes. Our results suggest that vaccine formulations, like Fluad, that specifically structure HA antigen into spiked nanodisc displays can offer examples of how to improve multivalency and immunogenicity of HA subunit vaccines to formulate more efficacious influenza vaccines for pandemic preparedness and universal influenza vaccine development. More broadly, these structure-guided approaches to influenza subunit vaccines containing HA can be applied to viral glycoproteins from other viruses with pandemic potential such as HIV, Ebola, and SARS coronaviruses.

## Materials and Methods

### Negative-Stain Electron Microscopy (EM) Specimen Preparation

Samples (e.g. Flucelvax, Fluzone HD, Flublok, and Fluad) were diluted in PBS to an HA concentration of 10 µg/mL. EM sample grids were prepared by applying 5 µL of the sample to a glow-discharged Formvar/carbon (F/C) 300 mesh grids. The specimens were stained with 3% phosphotungstic acid at pH 7 following previously reported methods ^38^. Data were acquired using a Tecnai 12 electron microscope (ThermoFisher) with a LaB_6_ filament operating at 100 kV and nominal magnifications of 67,000x, yielding a magnified pixel size of 1.8 Å/pixel. All images were recorded on a 4kx4k OneView camera (Gatan, Pleasaton, CA) using SerialEM software. Data for class averages was typically acquired in a single session lasting 1 to 2 hours, and yielding 100-200 micrographs ^39^. For 2018-2019 Flucelvax, 134 micrographs (12652 particles) were collected; for 2018-2019 Fluzone HD, 136 micrographs (15739 particles) were collected; for 2018-2019 Flublok, 27 micrographs (4575 particles) were collected; and for 2018-2019 Fluad, 97 micrographs (949 particles) were collected.

Negative stain EM data were processed using RELION3 software ^40^. The contrast transfer function (CTF) for each micrograph was estimated using CTFFIND4 software ^41^. For 2018-2019 Flucelvax, 500-1000 particles were manually picked. RELION3 2D classification was then used to generate auto-picking templates, which were used as references to repick particles on all micrographs, yielding a total of 12652 particles that were extracted at a box size of 256 pixels. These particles were aligned and classified using RELION3 2D Classification; the 2D averages were culled to remove poorly resolved classes (e.g. false positive picks or poorly aligned particles) from the particle set. Results from the final round of 2D classification were used to characterize the sample. Other negative stain datasets for Flublok, Fluad, and Fluzone were processed similarly.

### Monte Carlo Simulation

The crystal structure of HA in complex with stem Fab C179, PDB code 4HLZ, was used to model steric clashes between stem antibodies when bound to HA. Using the PERL scripting language, coordinates for the complex were rotated a random amount, then translated a random amount up to 25 nm. The software Chimera ^42^ was utilized directly, and called via PERL scripts to create the atomic coordinates in PDB format, and measure select distances and angles specified by a Chimera command file. Steric clashes between molecules were scored using software MolProbability Probe ^43^. A cycle of model generation, measurement, and clash scoring constituted one iteration of the Monte Carlo Method, and 500,000 such iterations were run. Plots were generated using the statistical software R ^44^.

### Cryo-Electron Tomography

Cryo-electron microscopy grids for 2018-2019 Flublok and 2018-2019 Fluad were prepared separately using similar procedures. Samples were mixed with 5 or 10 nm gold particles as fiducials. Using a Leica EM GP plunge freezer (Leica Microsystems), aliquots of 4 µL of sample were applied to glow-discharged, Quantifoil C Flat R2/2 200 mesh grids, blotted for 1.5 s in 99% humidity chamber, and plunge-frozen in liquid ethane. Flublok and Fluad tilt series were collected using Falcon II and Falcon 3 cameras on Titan Krios microscopes operating at 300 kV using a Volta phase plate (VPP) (FEI). Zero loss energy filtered data was collected with a 70 μm objective aperture at 3.3 Å pixel size (nominal magnification 26,000x) with a defocus range of −3.5 to −5.5 μm. Tilt series collected were acquired at 2.9 Å pixel size (nominal magnification 29,000x) with a −2.5 μm defocus. All single-axis tilt series data were collected over −/+ 60° with 2° tilt increments. Tilt series datasets were collected under low-dose conditions with a cumulative electron dose of 100 e^-^/Å^2^. Additional, Fluad tilt series data was collected with a Titan Krios microscope (ThermoFisher) operating at 300 kV with using similar data collection parameters reported above but with a GIF-2002 energy filter (slit width, 20 eV) coupled with a Gatan K2 camera (Gatan Inc.). Images within the tilt series were mutually aligned by using the gold particles as fiducial markers and the 3D volumes (tomograms) were reconstructed as implemented the IMOD and EMAN2 software packages ^45,46^.

Docking of HA trimers into subtomograms was performed using Chimera ^42^. 20 subtomograms of Flublok were analyzed, and 7 subtomograms of Fluad were analyzed. Scripted commands in chimera were used to measure distances and angles between HA trimers. PERL scripts were used to parse data in preparation for plotting, and R was used to generate plots.

For Subtomogram averages of Fluad discs, particles were picked from tomograms that had been reconstructed with EMAN2.91 ^46^. Particles were extracted and subtomogram averaged using RELION2.1 ^47^. To resolve the central lipid comprising the discs, particle box sizes was chosen to be 64 pixels, which was 346 Å at a working pixel size of 5.4 Å/pixel. To arrive at a single best-fit measurement of the disc thickness, D64 symmetry was imposed. 58 nanodiscs were used in the final subtomogram average structure.

### ELISA for Epitope Accessibility

For determination of available stem binding epitopes, vaccine formulations were each diluted to arrive at equivalent concentrations of H1 HA. Vaccine was applied to 96-well plates at 1.25 μg/mL and incubated overnight at 4°C. Plates were washed between all experimental step with PBS plus 0.1% Tween 20 (405 TS ELISA Plate Washer, Agilent Technologies). Blocking (1% Omniblok, AmericanBio, Inc., and 0.1% Tween 20 in PBS) occurred for 30mins at room temperature prior to application of the primary antibody (FI6v3, CR6261, and C179 provided by the Vaccine Research Center, Bethesda, MD) and incubated for 2 hr at room temperature. Plates were incubated for 1 hr at 37°C following the addition of an HRP conjugated secondary antibody (goat anti-human IgG (H+L) or goat anti-mouse IgG (H+L), (Thermo Fisher). Colorimetric detection occurred for 15 min at room temperature and absorbance was read at 405 nm (1-Step ABTS, Thermo Fisher). Samples were run in quadruplicate. Endpoint titer levels were statistically defined per plate ^48^. Statistical calculations were carried out with the software Prism (GraphPad).

For determination of mouse sera immunogenicity, antigen was applied to 96-well plates and incubated overnight at 4°C (1.25 μg/mL). Plates followed the same procedure as above with primary serum samples from vaccinated mice plated at an 8:1000 dilution with two-fold dilutions steps following. Statistical analysis was conducted for analysis of variance (ANOVA) using the F-test. The F statistic is reported (F=) with the parenthesis being the degrees of freedom within groups separated by a comma followed by the significance level (p) at the end.

### Mouse Immunization and Challenge

All mouse experiments were performed under protocols approved by the Animal Care and Use Committee (ACUC) at the National Institute of Allergy and Infectious Diseases (NIAID). 15 female BALB/c mice (Taconic Biosciences) aged 8-10 weeks were randomly assigned to a vaccine group. Groups of mice received one of five types of intramuscular injections for immunization on day 0 (prime) and day 21 (boost): saline, Fluad, Flublok, Fluzone, and Flucelvax. Mice were injected intramuscularly (50 μL/leg) at the manufacturers supplied concentration. Mice were weighed weekly to ensure health with bleeds occurring on day 0, day 14, and day 35 to track immunogenicity.

On day 42 mice were anesthetized and intranasal challenged with 25ul of influenza H1N1 A/Michigan/45/2015 at 10xMLD_50_ (50% Mouse Lethal Dose). The initial survival experiments used five mice per vaccine group and were observed twice daily for survival criteria (humane endpoints observed following a 30% decrease from initial body weight) until day 56, when all surviving mice were humanely euthanized. Differences in survival rates were compared using a Kaplan-Meier survival analysis (Graph Pad Prism). A replicate study was conducted with all mice being euthanized 3 days post challenge for the collection of lung tissue for pathology. A control mouse group for pathology studies received “vaccinations” with saline on day 0 and day 21, and “challenge” with saline supplemented with 0.1% bovine serum albumin (BSS-BSA) on day 42.

### Determination of Influenza H1N1 A/Michigan/45/2015 Mouse Lung Titer

TCID50 assays were conducted by infecting MDCK-ATL cells with 1:5 serial dilutions of each mouse lung homogenate in duplicate in serum-free MEM with 1 μl/ml TPCK-Trypsin and incubated at 37°C and 5% CO_2_ for 96hrs. Cytopathic effect (CPE) was determined following incubation by staining with crystal violet and TCID50 was calculated using the Reed-Muench method ^49^.

### Immunohistochemistry

Formalin-fixed paraffin-embedded mouse lung sections were labeled for immunohistochemical staining using a rabbit polyclonal anti-Influenza A virus nucleoprotein antibody, CAT# GTX125989 (GeneTex). Staining was carried out on the Bond RX (Leica Biosystems) platform according to manufacturer-supplied protocols. Briefly, 5 µm-thick sections were deparaffinized and rehydrated. Heat-induced epitope retrieval (HIER) was performed using Epitope Retrieval Solution 2, pH 9.0, heated to 100°C for 20 minutes. The specimen was then incubated with hydrogen peroxide to quench endogenous peroxidase activity prior to applying the primary antibody at a dilution of 1:4000. Detection with DAB chromogen was completed using the Bond Polymer Refine Detection kit (Leica Biosystems). Slides were finally cleared through gradient alcohol and xylene washes prior to mounting and coverslipping. Sections were examined by light microscopy using an Olympus BX51 microscope and photomicrographs were taken using an Olympus DP73 camera. An unbiased blinded then unblinded approach was used to analyze the of immunization conditions of mice using commercial influenza vaccines with histopathology after H1N1 challenge.

### Data deposition

Tomographic reconstructions have been deposited in the Electron Microscopy Data Bank under accession numbers EMDB-27232 (Fluad spiked nanodisc) and EMD-27233 (Flublok starfish).

## Acknowledgements

This work was supported by the Intramural Research Program of the National Institute of Allergy and Infectious Diseases. We thank Jeffery Taubenberger and Jae-keun Park for kindly providing the headless trimeric stem protein and the International Reagent Resource for HA proteins. We thank Masaru Kanekiyo and Barney Graham for providing stem antibodies. We also thank Vinod Nair, Cindi Schwartz, Elizabeth Fischer, and Rick Huang for help in cryo-EM data collection. This work utilized the computational resources of the NIH HPC Biowulf cluster (http://hpc.nih.gov), and the Office of Cyber Infrastructure and Computational Biology (OCICB) High Performance Computing (HPC) cluster at the National Institute of Allergy and Infectious Diseases (NIAID), Bethesda, MD

