## Supplemental Material for "Commercial influenza vaccines vary in both the structural arrangements of HA complexes and in induction of antibodies to cross-reactive HA epitopes"

| Name | Vaccine Type | Valency | Manufacturer | Total HA | HA/Strain | H1N1 Strain | H3N2 Strain |
|---|---|---|---|---|---|---|---|
| Fluad | Subunit, Adjuvanted | Trivalent | Seqirus | 45ug/0.5ml | 15ug/0.5ml | A/Singapore/GP1908/ 2015 IVR-180 | A/Singapore/INFIMH-16-2016 IVR-86 |
| Flublok | Recombinant | Quadrivalent | Sanofi Pasteur | 180ug/0.5ml | 45ug/0.5ml | A/Michigan/45/2015 | A/Singapore/INFIMH-16-2016 |
| Fluzone HD | Split Virus | Trivalent | Sanofi Pasteur | 180ug/0.5ml | 60ug/0.5ml | A/Michigan/45/2015 x-275 | A/Singapore/INFIMH-16-2016 IVR-86 |
| Flucelvax | Subunit | Quadrivalent | Seqirus | 60ug/0.5ml | 15ug/0.5ml | A/Singapore/GP1908/ 2015 IVR-180 | A/NorthCarolina/04/2016 |

| Name | Influenza B- Victoria | Influenza B- Yamagata | Inactivation | Method of Disruption | Reported Purification Method |
|---|---|---|---|---|---|
| Fluad | B/Maryland/15/2016 | | Formaldehyde | Cetyltrimethyl ammonium bromide | Zonal centrifugation |
| Flublok | B/Maryland/15/2016 | B/Phuket/3073/2013 | | Triton X-100 | Column chromatography |
| Fluzone HD | B/Maryland/15/2016 BX-69A | | Formaldehyde | Octyphenoleth oxylate (Triton X-100) | Linear sucrose density gradient solution using continuous flow centrifuge |
| Flucelvax | B/Iowa/06/2017 | B/Singapore/INFTT-16-0610/2016 | β-propiolactone | Cetyltrimethyl ammonium bromide | "several process steps" |

**Figure S1. Commercial influenza vaccines used in this study**. Details are provided for the 4 commercial vaccines studied: Flucelvax, Flublok, Fluzone and Fluad. Both the total HA concentration and the concentration of HA components such as H1 or H3 HA subtypes is denoted.

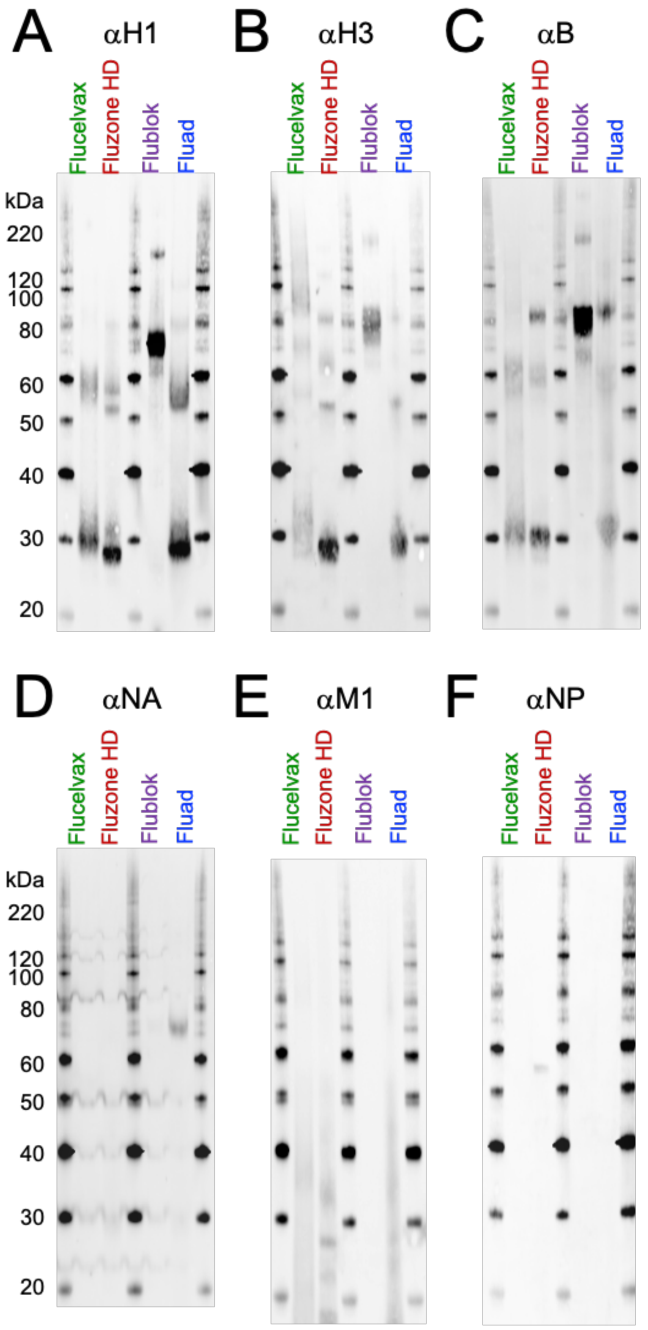


**Figure S2. Analyzing the composition of commercial influenza vaccines for influenza structural proteins HA, NA, M1 and NP.** (A-F) Reactivity analysis via Western blot of commercial influenza vaccines (Flucelvax, Fluzone, Flublok, Fluad) with primary antibodies to hemagglutinins (A) H1, (B) H3, and (C) influenza B. Also probed were (D) neuraminidase (NA), (E) matrix M1, and (F) nucleoprotein (NP). Molecular weight standards are given in panels A and D, and are the same for all panels.

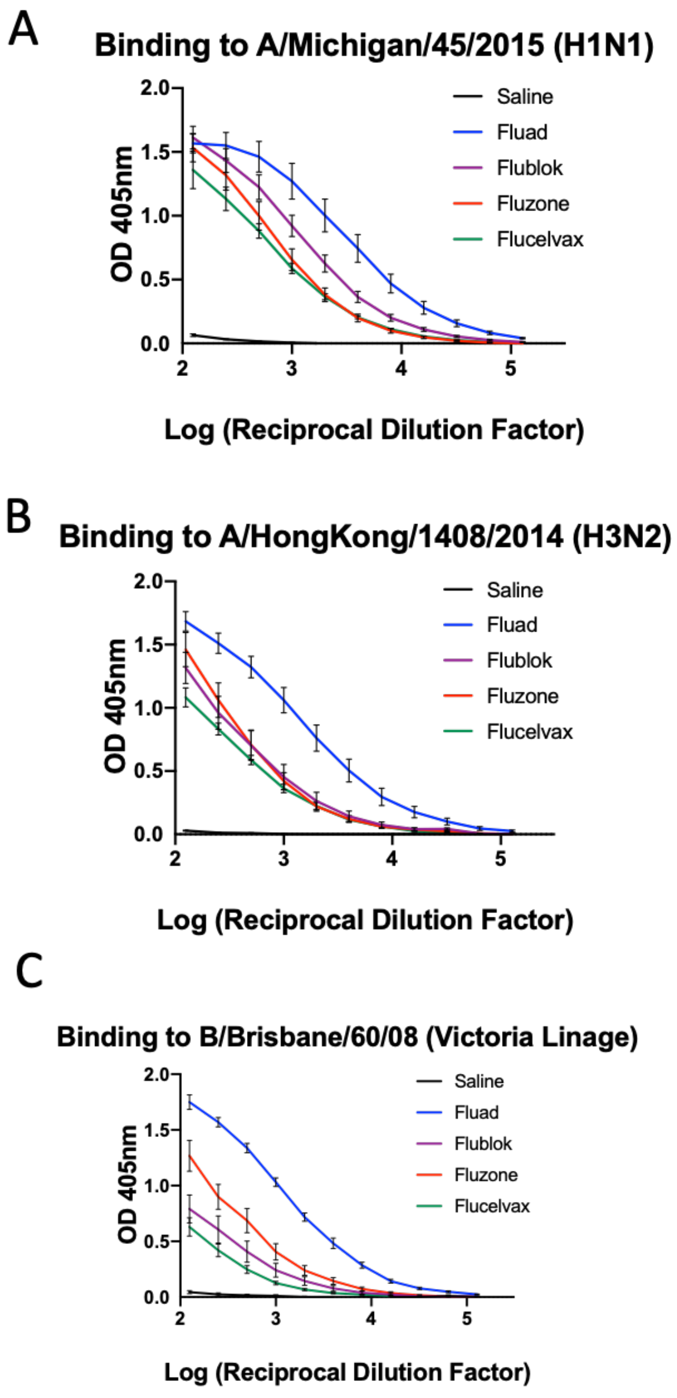


**Figure S3. Homosubtypic binding to hemagglutinin proteins from vaccinated mouse sera**. Serially diluted sera from mice immunized with commercial vaccines Fluad, Flublok, Fluzone and Flucelvax were analyzed by ELISA for binding to antigenically matched HA proteins representing (A) H1N1, (B) H3N2, and (C) B-Victoria lineage influenza viruses.

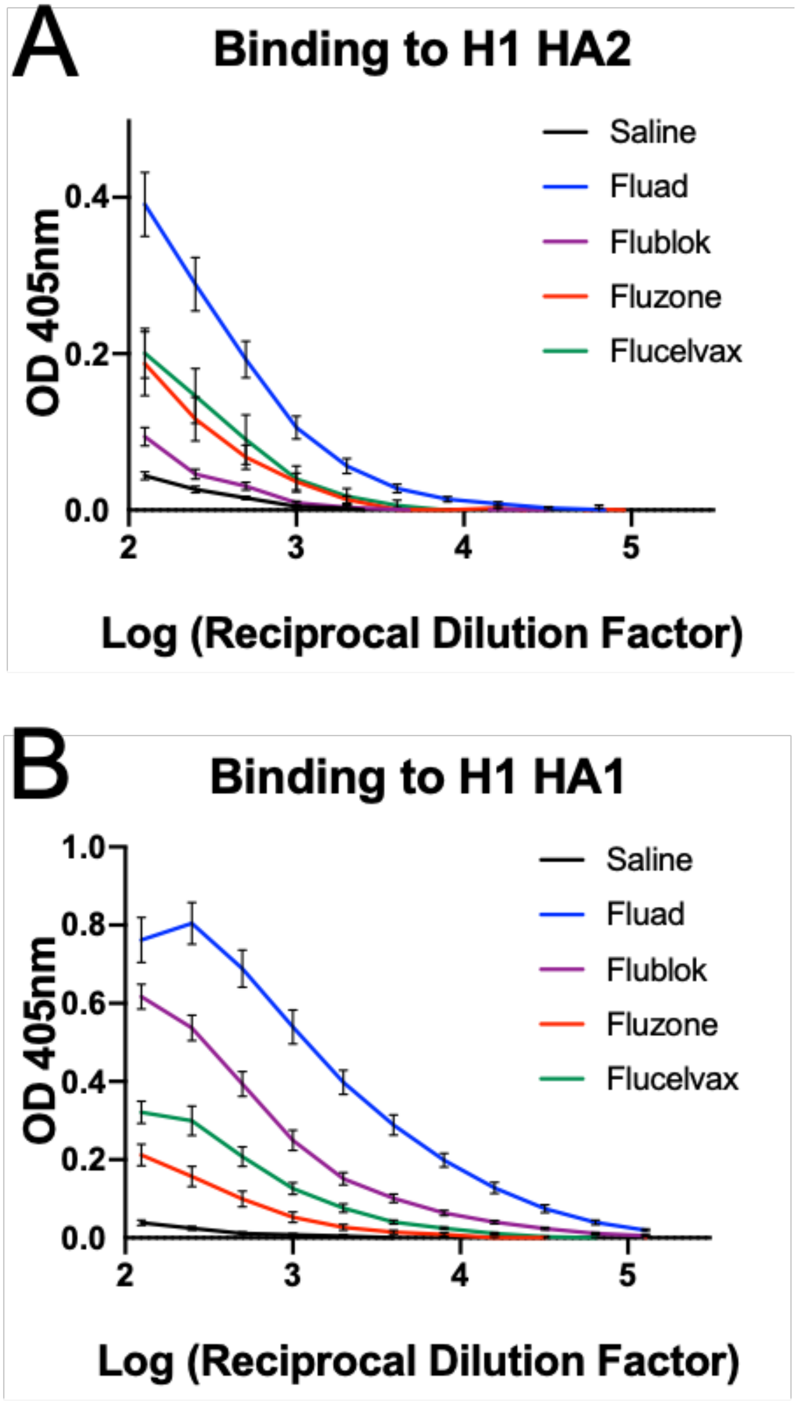


**Figure S4. HA head and stem epitope targeting of elicited antibodies from commercial influenza vaccines.** Binding to recombinant protein representing the (A) recombinant headless H1 HA2 stem construct and (B) recombinant H1 HA1 head was compared by ELISA using serially diluted sera from mice immunized with commercial vaccines Fluad, Flublok, Fluzone, Flucelvax, and with saline as a control. Curves were used to derive endpoint titers shown in main-text figure 5.

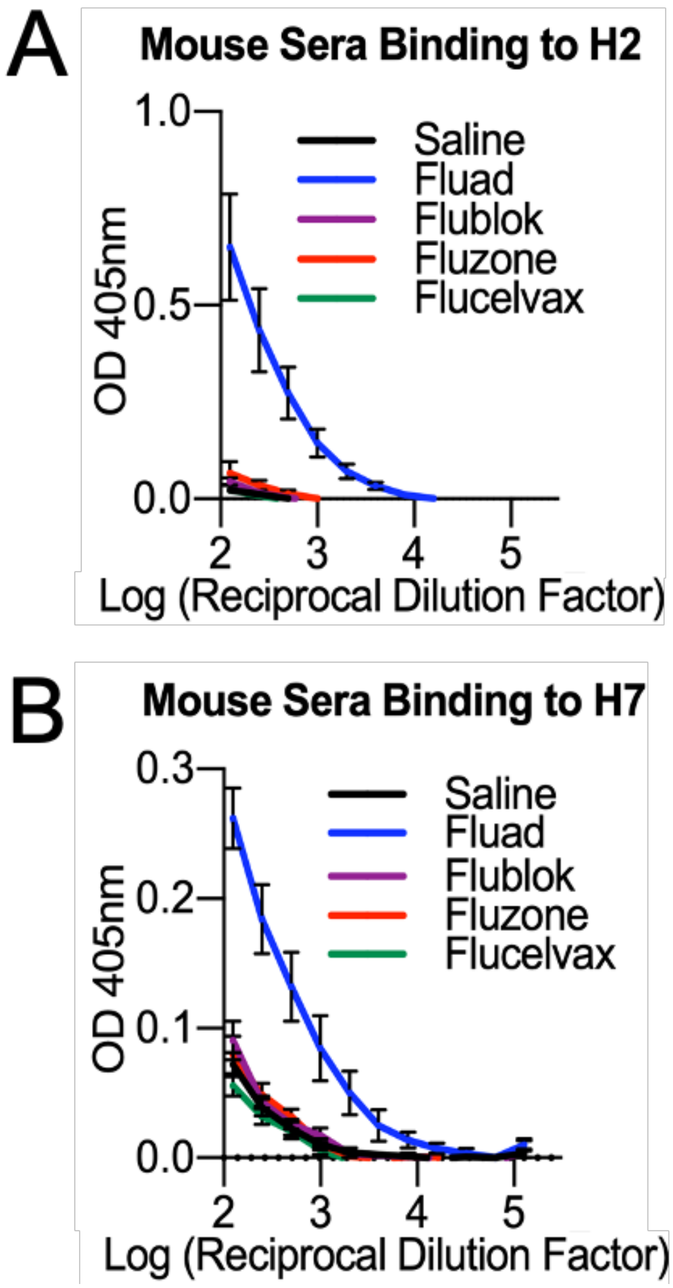


**Figure S5. Cross-reactive immunogenicity of vaccinated mouse sera for H2 and H7 HA.** Binding to full-length (A) H2 and (B) H7 proteins were compared by ELISA of serially diluted sera from mice immunized with commercial vaccines Fluad, Flublok, Fluzone, Flucelvax, and with saline as a control. Curves were used to derive endpoint titers shown in main-text figure 5.

a =airway
I =pulmonary
interstitium

**A** Mouse IDs/ Vaccine immunization condition

| ID | Condition | Day |
|---|---|---|
| 671 | BSS BSA | 3 |
| 672 | BSS BSA | 3 |
| 646 | Saline | 3 |
| 647 | Saline | 3 |
| 651 | Flucelvax | 3 |
| 652 | Flucelvax | 3 |
| 656 | Fluzone HD | 3 |
| 657 | Fluzone HD | 3 |
| 661 | Flublok | 3 |
| 662 | Flublok | 3 |
| 666 | Fluad | 3 |
| 667 | Fluad | 3 |
| 673 | BSS BSA | 6 |
| 674 | BSS BSA | 6 |
| 648 | Saline | 6 |
| 649 | Saline | 6 |
| 653 | Flucelvax | 6 |
| 654 | Flucelvax | 6 |
| 658 | Fluzone HD | 6 |
| 659 | Fluzone HD | 6 |
| 663 | Flublok | 6 |
| 664 | Flublok | 6 |
| 668 | Fluad | 6 |
| 669 | Fluad | 6 |
| 675 | BSS BSA | 14 |
| 655 | Flucelvax | 14 |
| 660 | Fluzone HD | 14 |
| 665 | Flublok | 14 |
| 670 | Fluad | 14 |

**B** Blinded evaluation of lung pathology

| Viral Staining | TCL |
|---|---|
| Neg | Neg |
| Neg | Neg |
| Interstitial | Med |
| Interstitial | High |
| Ariway | Low |
| Ariway | Low |
| Ariway | Low |
| Ariway | Low |
| Neg | Neg |
| Ariway | Low |
| Neg | Neg |
| Neg | Neg |
| Neg | Neg |
| Neg | Neg |
| Interstitial | Med |
| Interstitial | High |
| Ariway | Low |
| Ariway | Low |
| Ariway | Low |
| Ariway | Med |
| Neg | Neg |
| Ariway | Low |
| Neg | Neg |
| Neg | Neg |
| Neg | Neg |
| Ariway | Low |
| Ariway | Low |
| Ariway | Low |
| Ariway | Low |

**C**

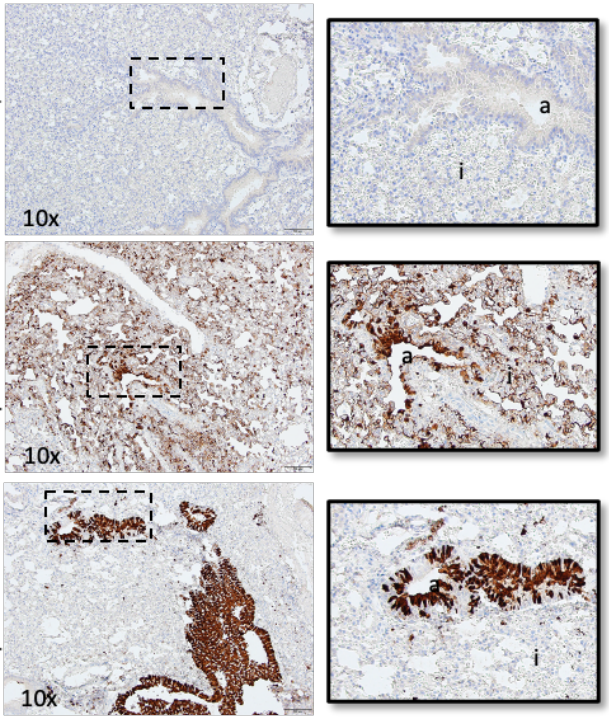


**Figure S6. Analysis approach to histopathology after H1N1 challenge**. An unbiased, blinded method was employed for analysis of histopathology of mouse lungs following H1N1 challenge. (A) Mouse identification numbers (IDs) were generated to blind the type of immunogen and challenge given. (B) Blinded evaluation of tissue samples was used to classify viral presences as negative, low, medium, and high. (C) Selected sample images illustrate immunohistochemistry staining (IHC) against the influenza nucleoprotein protein for groups corresponding to negative healthy lung, interstitial antigen staining, and airway antigen staining.

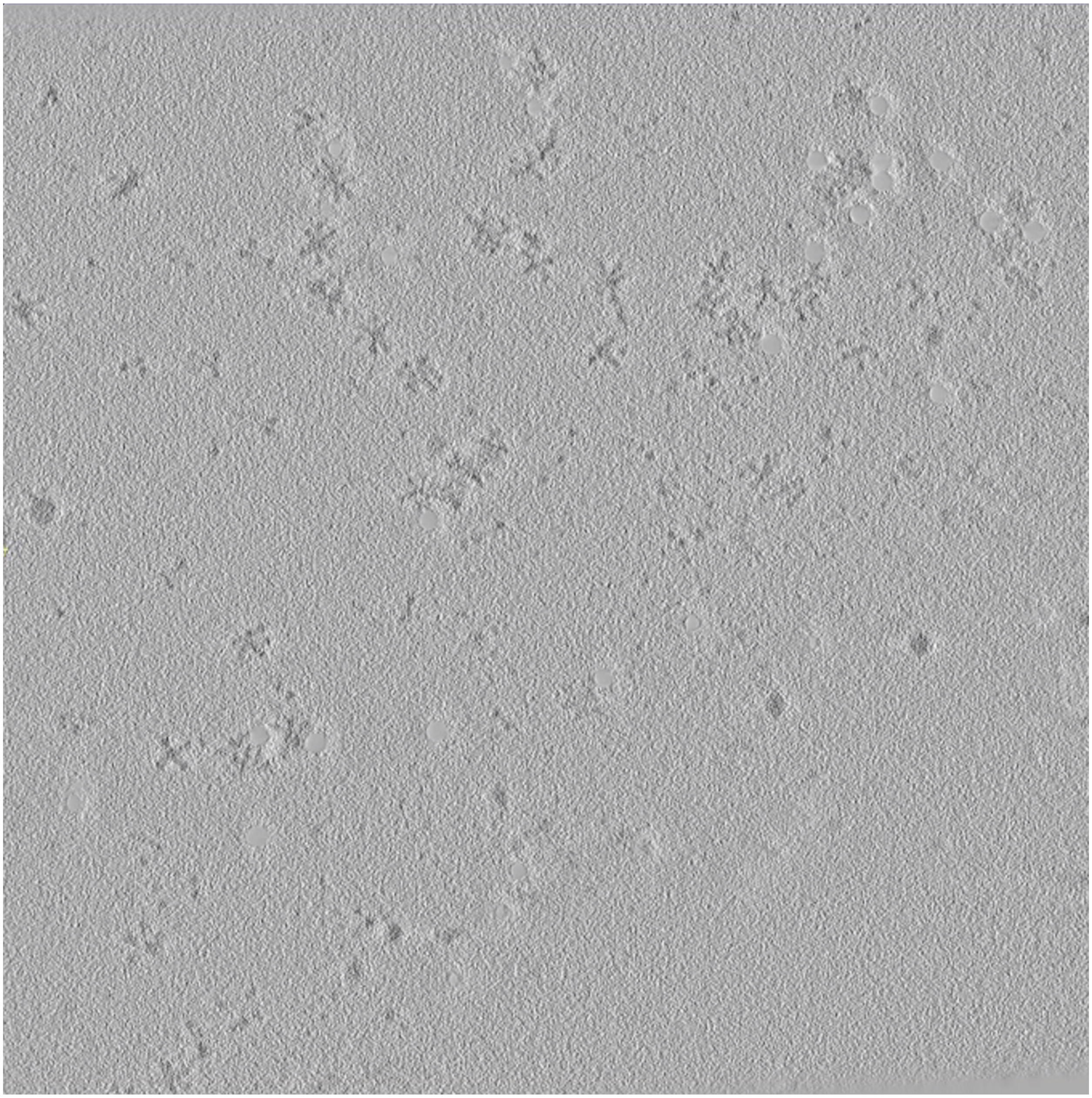

**Movie S1. Slicing through a 3D tomogram of the commercial influenza virus vaccine Flublok.** HA complexes appear as starfish-like structures. Within each complex, HA trimers emanate outward from a central focus in all directions.

.

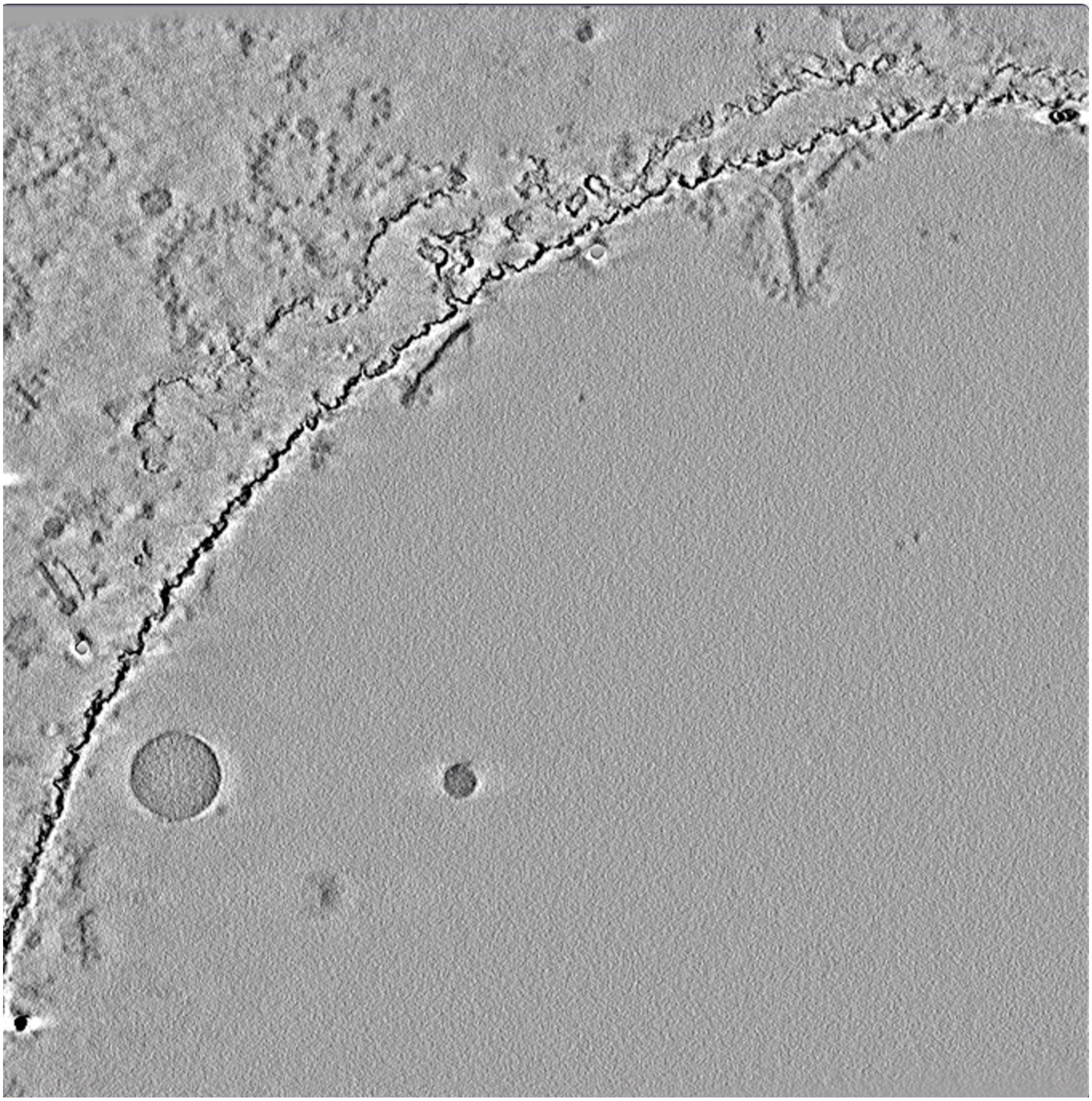

**Movie S2. Slicing through a 3D tomogram of the commercial influenza virus vaccine Fluad**. Two Fluad spiked nanodiscs are shown proximal to the carbon edge of the hole on the EM grid. As the visualization plane slices through the sample, a dark band of density transits from one side of the disc to the other, which is the result of the oblique angle of the complex to the imaging plane. One Fluad spiked nanodisc is in the upper right, while another Fluad spiked nanodisc is to the left. An example of an adjuvant vesicle is in the lower left and appears as an approximate grey sphere.

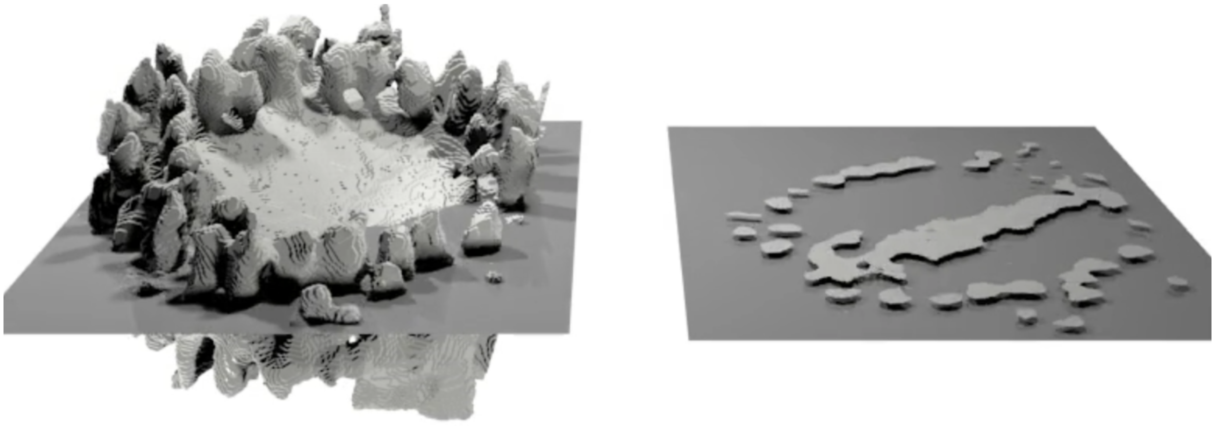

**Movie S3. Fluad spiked nanodisc from cryo-electron tomography viewed as a 3D solid surface rendered as slices of the volume. (**Left) Successive plane slicing through a solid 3D surface rendering of the 3D tomographic volume of a Fluad complex (i.e. spiked nanodisc). (Right) Corresponding planes only showing the density in the plane of the slice. The glycoprotein spikes of the complex appear as dotted densities around a central band of membrane.
